# The emergence of cooperation by evolutionary generalization

**DOI:** 10.1101/2021.01.27.428436

**Authors:** Félix Geoffroy, Jean-Baptiste André

## Abstract

In principle, any cooperative behaviour can be evolutionarily stable as long as it is incentivized by a reward from the beneficiary, a mechanism that has been called reciprocal cooperation. However, what makes this mechanism so powerful also has an evolutionary downside. Reciprocal cooperation faces a chicken-and-egg problem of the same kind as communication: it requires two functions to evolve at the same time –cooperation and response to cooperation. As a result, it can only emerge if one side first evolves for another reason, and is then recycled into a reciprocal function. Developping an evolutionary model in which we make use of machine learning techniques, we show that this occurs if the fact to cooperate and reward others’ cooperation become general abilities that extend beyond the set of contexts for which they have initially been selected. Drawing on an evolutionary analogy with the concept of generalization, we identify the conditions necessary for this to happen. This allows us to understand the peculiar distribution of reciprocal cooperation in the wild, virtually absent in most species –or limited to situations where individuals have partially overlapping interests, but pervasive in the human species

---

As a general rule, natural selection does not favour strategies that support the common good. Except for genetically related organisms, any action that entails a fitness cost is counterselected, no matter how beneficial it is for other individuals (1–3). In consequence, natural selection results in a “cooperation load”, a set of cooperation opportunities that are missed by evolution.

There is, however, a solution that can make, at least in principle, any cooperative behaviour adaptive as long as it has a net beneficial effect. It needs to be incentivized with a reward (4–17). If an individual has the opportunity to pay a personal cost *c* to bring a benefit *b* to another, the latter can encourage cooperation by promising to provide a reward *r* in exchange. This is adaptive for both individuals as long as *r* is sufficient to compensate for the cost of cooperation, but not to the point of outweighing its benefit –that is, as long as *c <r <b*. Hence, any cooperative action with a net social benefit (*b > c*) can be enforced with an appropriate reward. In the economic theory of incentives, rewards are usually assumed to be part of a contract enforced by institutions (Laffont and Martimort 2001) but non-cooperative game theory shows that they can also be made evolutionarily stable endogenously if interactions are repeated. In this framework, rewarding others is evolutionarily stable because it allows to gain access to similar social interactions in the future, with the same or with other partners. In the following, we will refer to this mechanism as “reciprocal cooperation”, in a broad sense that includes all mechanisms whereby cooperation is incentivized by a reward.

There is one problem with this mechanism, however. Unlike rational individuals who can foresee the consequences of their decisions in the future, biological evolution is shortsighted. Every genetic mutation involved in a biological function must be advantageous when it appears. As a consequence, the evolution of reciprocal cooperation raises a bootstrapping problem (18, 19). Cooperation cannot be adaptive in the absence of rewards because it is costly; and rewards are not adaptive unless a significant fraction of individuals are already willing to cooperate conditionally on the presence of a reward. Hence, the evolution of reciprocal cooperation would require that the two functions appear together at once, which is highly unlikely.

Communication is another type of biological interaction that is notoriously subject to the same type of bootstrapping problem. Two complementary traits must be present for communication to occur, a signal and a response, neither of them being adaptive without the other. For this reason, communication cannot evolve from scratch. Evolutionary biologists, however, have long understood how evolution can solve this problem. Communication can evolve if one side –the signal or the response– first evolves for another reason, and is then recycled into a communicative function (20–22).

The solution must be along the same lines in the case of reciprocal cooperation. Reciprocal cooperation can evolve if one side of the problem evolves first for an independent reason and is then recycled into a reciprocal cooperation function (23, 24). Understanding the emergence of reciprocity, and the possible constraints that limit its scope in the living world, therefore involves understanding what makes possible, and also what limits, the recycling of a biological function in a different context than it has initially been selected for. This is what we are aiming to do in the present article.

Leveraging the performance of cognitive devices outside the range of contexts they have originally been made for is at the heart of *machine learning*. The statistical theory that forms the basis of this technique shows that the ability of machines to *generalize* is not a contingent, lucky, outcome of learning. On the contrary, generalization obeys a systematic logic, and occurs in specific circumstances that can be understood (25).

Because natural selection is an adaptive process, like learning, the concept of generalization can readily be extended to biological evolution (26–35). In this paper, we will show that reciprocal cooperation can evolve for the same reason that machines generalize. Reciprocal cooperation evolves if the fact to cooperate and reward others’ cooperation become general abilities that extend beyond the very set of contexts for which they have initially been selected, and the conditions necessary for such evolutionary generalization to take place can be understood with machine learning concepts. We will show that this provides key insights on the nature of reciprocal cooperation in the wild, its limits in extant species, and its phylogenetic distribution.

## Games of life

Evolutionary game theoreticians generally study the evolution of social behaviours in a single game, isolated from all other facets of individuals’ life. This entails the implicit assumption that individuals evolve separate rules of decision adapted to each and every game, which poorly captures how individuals make decisions in practice. An individual’s life comprises a variety of situations and, although each situation entails a specific problem, the individual must eventually make all decisions with the same cognitive device. Hence, models need to capture the fact that the decision machinery of individuals is selected to deal in the same time with a set of games instead of just one (36–39). This has major implications, we will see, for the evolution of cooperation.

In our model, individuals are confronted, across their life, to a set of situations, chosen in a random order, which we each call a *game*. Each game involves two individuals, a truster and a trustee, both characterized by a decision mechanism modeled as an artificial neural network, whose connection weights evolve by natural selection (see *Methods* for details). For simplicity, we assume that the truster and the trustee are drawn from two separate populations, but the model would be almost identical otherwise. In each game, the truster first decides whether he wants to invest or decline the game. If he decides to invest, he pays a personal cost *c* while the trustee gets a benefit *b*. The trustee then has the opportunity to decide on a quantitative reward *r* that she can transfer to the truster in return. There are four types of games depending on the values of *b* and *c*.

In **Selfish** games, the truster personally benefits from investing (that is, *c* < 0) and the trustee may either benefit or be unaffected by the game (that is, *b* ≥ 0). Selfish games capture both situations of common interest, also called byproduct cooperation (40), where the truster and the trustee benefit from the same action, and also the very large amount of situations where the truster simply acts in a self-serving way without involving the trustee.

In **Wasteful** games, the investment is costly to the truster (*c >* 0) and it does not benefit the trustee sufficiently to offset this cost (*b < c*). Wasteful games represent the very frequent situations where no mutually beneficial interaction can be found between players.

In **Cooperative** games, the investment is costly to the truster (*c >* 0) and generates a benefit *b > c* for the trustee. Therefore, the truster has no immediate benefit in cooperating but a mutually beneficial agreement can be found if the trustee rewards with *r* ∈]*c,b*[. Cooperative games comprise all situations where cooperation has a net social benefit but raises an evolutionary bootstrapping problem. Throughout the rest of the paper, for the sake of simplicity we will speak of “cooperation” to mean “investing in a Cooperative game” (in a somewhat unorthodox use of the term, see 3). Our aim is to understand how cooperation, in this sense, can evolve.

Finally, we consider a fourth category of games, that we call **Interdependent**, capturing situations where (i) the investment benefits the trustee, and (ii) the eventual net benefit for the truster depends on the trustee’s quality, that is situations of *partner-dependent common interest* (40–46). For instance, when an ant colony invests in protecting an acacia tree the eventual benefit of the investment depends on the acacia’s actual value as a shelter (47). When a pollinator visits a flower, the time spent in the flower can be worth it or not, depending on flower’s quality (48). When an individual spends time and energy in a collective action with conspecifics (e.g., performing a collective hunt, inspecting a predator, etc.), the eventual benefit of the investment depends on the quality of the partner(s) (e.g. 49). Etc. Formally, Interdependent games are characterized by a positive cost for the truster (*c >* 0), a benefit *b* for the trustee, and what we call an “automatic” reward *r_a_*, whose value depends on the trustee’s quality. In practice, in our model, each time two players play an Interdependent game, the value of the automatic reward is simply drawn at random among two possible values, *r*_*a*1_ = 0 or *r*_*a*2_ > *c*, each with a 0.5 probability. Finally, without loss of generality, we assume that *b > c* in all Interdependent games such that a mutually beneficial agreement can always be found.

In all types of games, beyond the possible automatic reward, the trustee has the possibility to invest into an *active* reward, *r*, in order to provide a supplementary benefit to the truster. For simplicity, we assume that this active reward is a conservative transfer of fitness units, with a benefit *r* for the truster at the exact same cost for the trustee.

## Generalization creates latent rewards

We assume that, initially, before natural selection acts on the behaviour of individuals, the trustees do not actively reward any game (that is, *r* = 0 in all cases). As a result, the trusters are under selection to decline all Cooperative games, and the trustees’ propensity to reward these games is hence not under selection as the trusters never invest in them anyway (see Fig. 1 and *Methods*). This is the essence of the bootstrapping problem of cooperation. The key to solving this problem, however, lies in the existence of *latent* rewards, that is rewards that the trustees *would* offer in Cooperative games, if the trusters did play them. The evolution, or non-evolution, of cooperation depends critically on the existence and value of latent rewards. If the latent reward happens to be *r > c* in a given game, then cooperation can evolve in this game, as this reward constitutes a yet undiscovered benefit that the trusters are under selection to collect.

**Fig. 1.**
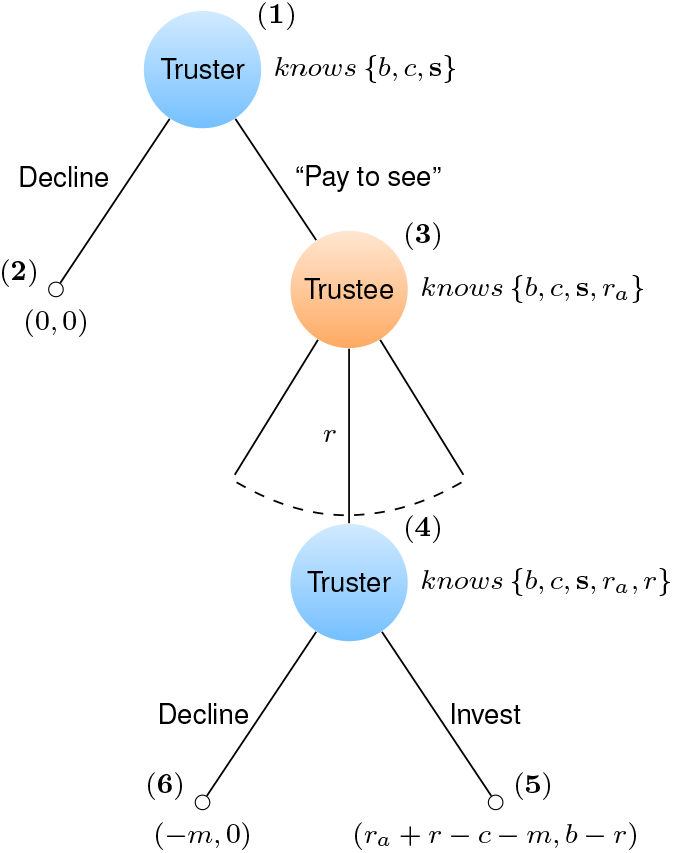
Outline of a game. (1) The truster is informed of all the game parameters that do not depend on the trustee and decides either to decline the game or to “pay to see” the trustee’s quality. (2) If the truster declines, the game ends with no payoffs and the next game is randomly chosen. (3) If the truster decides to “pay to see”, he pays a small monitoring cost, *m*, to find out about the total reward (*r_a_* + *r*) he would eventually get would he decide to play the game. This is meant to capture all the diverse manners in which one can obtain information on one’s partner. For instance, the truster could collect information by observing some aspects of the trustee’s phenotype that are correlated with her quality, by playing the game once to “test” the trustee, by finding out about the trustee’s past behaviour, etc. (4) At the last stage, we assume that the truster decides to (5) proceed with the game if *r* + *r_a_ > c* or (6) to decline if not.

The problem is that evolutionary game theory cannot directly tell us anything about latent rewards as they are, precisely, not under selection. Their evolution can nevertheless be understood provided one takes into account the fact that selection on *actual* rewards indirectly affects the evolution of *latent* ones. To understand this point, it is useful to draw up a parallel with machine learning. In machine learning, a machine (e.g. an artificial neural network), is trained to map a given set of inputs, called the training set, to specific outputs. But the aim of machine learning is not only to produce machines that can produce correct answers in contexts they have already encountered. The raison d’être of this technique comes from the fact that machines are able to *generalize* beyond their training set. That is, machines can also produce correct decisions in situations they have never been trained to deal with. And this ability to generalize is not a contingent, lucky, outcome of learning. On the contrary, generalization obeys a systematic logic, and occurs in specific circumstances that can be understood.

Being two adaptive processes, learning and evolution by natural selection have similar properties, and generalization therefore has an evolutionary counterpart (32). The set of situations for which a population is, or has been, under selection corresponds to the population’s training set, that is the situations for which the population received feedback from the environment via natural selection. Conversely, the set of situations for which a population has never been under selection, because they do not occur frequently enough to constitute a selective pressure, corresponds to the population’s test set. Any response on the test set is an unintended property of the way organisms generalize beyond their training set.

In our model, the trustees’ training set consists of the set of games in which the trusters do invest. Throughout their life, the trustees are confronted to these games and must decide on a reward. Hence, as a population, they receive feedbacks from natural selection. The trustees’ test set, on the other hand, consists of the set of games in which the trusters never invest. The trustees never actually have to decide on a reward in these games. Hence, they never receive any feedback from natural selection. Put differently, the trustees’ test set consists of all the games in which their ability (or inability) to reward trusters’ investments is only *latent*. Since the evolution of cooperation depends on the values of these latent rewards, it depends on the ability of the trustees to generalize.

To study the evolution of latent rewards in our multi-game framework, we first let the trustees evolve while keeping the trusters’ behavior as an exogenous constant. We assumed that the trusters always invested into Selfish and Interdependent games and always declined Wasteful and Cooperative games. Figure 2 shows the evolution of the trustees’ behaviour over time. In equilibrium, the trustees never rewarded Selfish games, as rewarding these games was superfluous, and they rewarded Interdependent games by adding an active reward *r > c − r_a_* when the automatic reward happened to be insufficient to compensate for the trusters’ cost (that is, when *r_a_ < c*), (Figure 2 solid lines). Moreover, as a side effect of these adaptive rewards, the trustees also had *latent* rewards: they rewarded the trusters in a fraction of Cooperative games even though they never actually encountered these situations (Figure 2 dashed line).

**Fig. 2.**
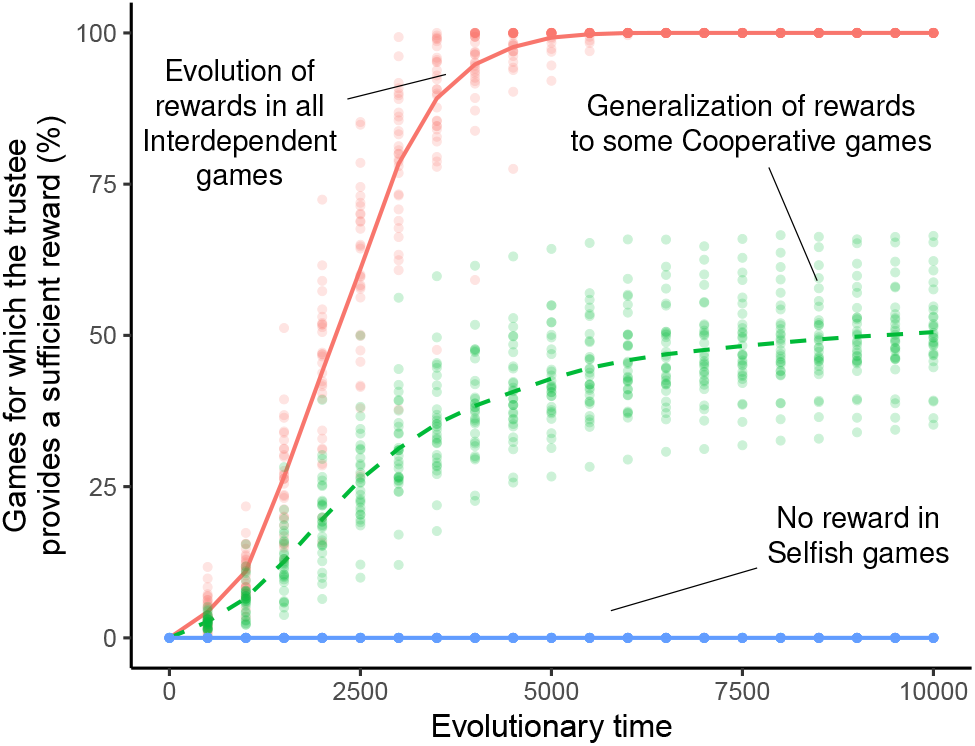
Adaptation and generalization in the trustees. On the y-axis, we indicate the percentage of games of each type for which the trustees provide a sufficient reward to induce the trusters’ investment. The solid lines show the adaptation of the trustees’ reward in Interdependent and Selfish games, through evolutionary time. The dashed green line shows the repercussions of this adaptation on the latent response of the trustees in Cooperative games –for which the trustees is not subject to selection. Wasteful games are not shown here (but see SI Figure S1). There are 2^5^ Interdependent games, *N_P_* = 1 input per payoff parameter, and *N_S_* = 50 spurious features. Mean over 30 simulations. See *Methods* for details on other parameters. Note that, for Selfish games, the truster always has an incentive to invest regardless of the trustee’s active reward. So here, we simply show the mean active reward provided in Selfish games (blue line), which is always zero.

Figure 3 shows how this generalization depended upon circumstances. In particular, it depended critically on the prevalence of Interdependent interactions in individuals’ ecology. When the environment contained very few Interdependent games, the trustees almost never generalized at the evolutionary equilibrium (Figure 3a). They rewarded optimally when in one of the few Interdependent games but under no other circumstances. Generalization was more significant, on the other hand, when many Interdependent games were present in individuals’ ecology. The trustees did not only reward optimally in Interdependent games, they were also able to calculate the right reward they should offer in some, or even sometimes in all, Cooperative games, even though they had never been selected to do so (Figure 3a).

**Fig. 3.**
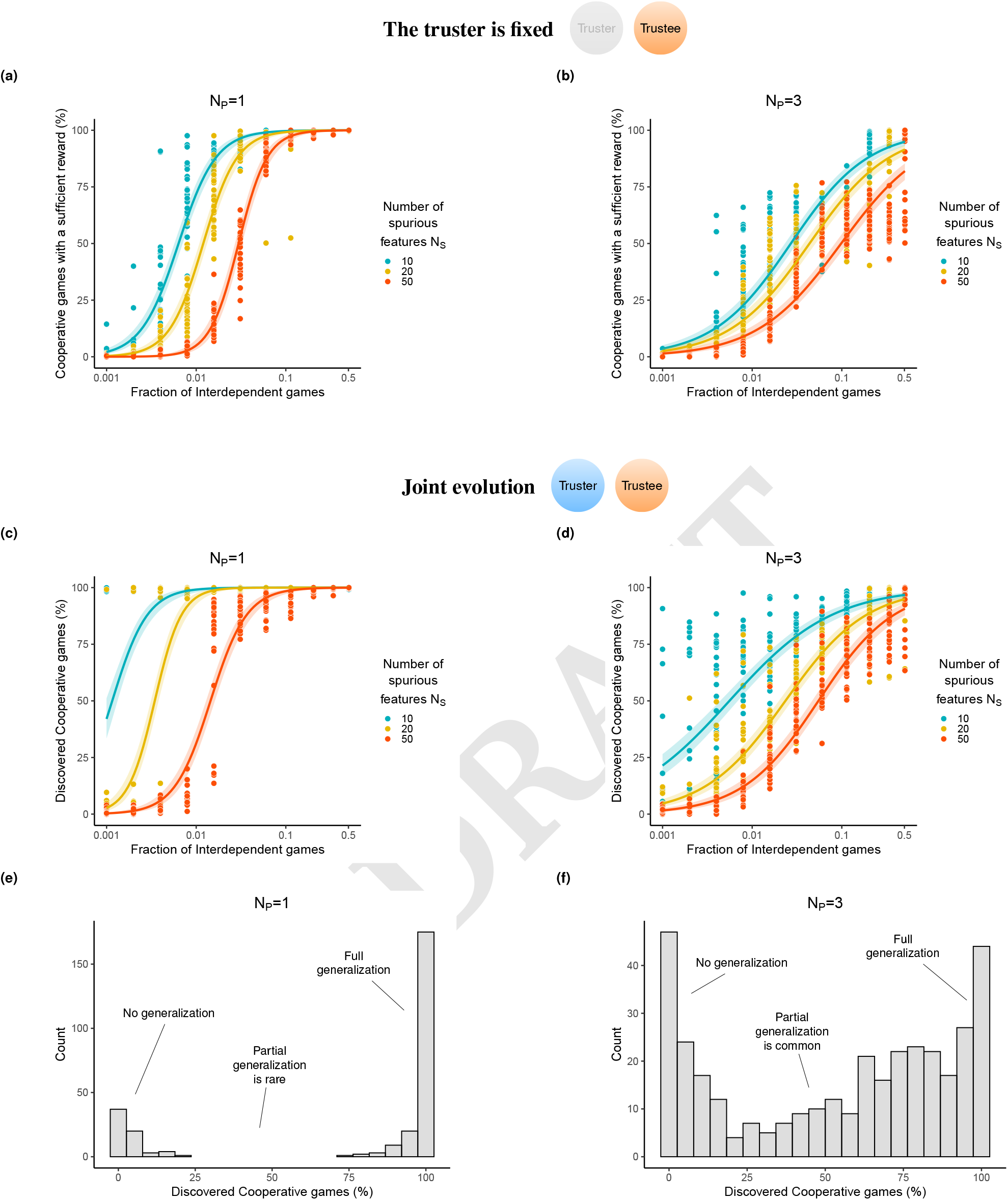
The extent of generalization in function of individuals’ ecology. **(a,b)** We assume here that only the trustees are evolving, i.e. the truster’s behaviour is fixed (see main text). We show the percentage of Cooperative games for which the trustees provide, in evolutionary equilibrium, a latent reward that would be sufficient to incentivize the trusters’ investment. Each point is a simulation result (30 simulations have been performed per parameter set). The solid line and shaded area correspond to the best logistic fit (mean and standard error). Results are shown for various numbers of spurious and genuine features (*N_S_* and *N_P_*, see *Methods*). **(c,d)** We assume here that the trusters and the trustees jointly evolve. We show the percentage of Cooperative games which are both played by the trusters and rewarded by the trustees in evolutionary equilibrium. Conventions and parameters are the same as in **(a)** and **(b)**. **(e, f)** Histograms showing the distribution of the percentage of “discovered” Cooperative games over all the simulation results shown, respectively, in **(c)** and **(d)**. See *Methods* for details on other parameters.

The reason why a small number of Interdependent games hinder generalization is the same as in the well-known problem of over-fitting with imbalanced data in machine learning. It is known that in classification tasks, if the training set is imbalanced (e.g. it includes many more negative than positive examples, or vice versa) then generalization is hardly possible (50–54). More precisely, if the number of positive examples in the training set is very small, the ANN typically learns to classify them accurately, but too specifically, reducing its classification performance in the test set, a situation called over-fitting.

Beyond the prevalence of interdependence in individuals’ ecology, the possibility of generalization also depended on the cognitive complexity of the task. We captured this complexity with two parameters, *N_P_* and *N_S_*, which represented respectively the number of sensory inputs that individuals had to integrate to evaluate the payoffs of a game, and the number of distracting inputs that could be spuriously correlated with the game payoff but did not allow for generalization (see Methods). As is standard in machine learning, the larger was *N_P_* and/or *N_S_*, the larger was the number of Interdependent games that were required to achieve full generalization (Figs. 3a and 3b).

## Latent rewards foster even more generalization

In addition to the evolution of the trustees, we then considered the fact that the trusters are also shaped by natural selection (see *Methods*). Generalization –or an equivalent process (see Fig. 4)– could then occur on both sides, reinforcing each other, and leading to the eventual discovery of more Cooperative games. The joint evolution of cooperation and reward in this case is schematized in Figure 4 and an example is shown in Figure 5. In comparison with the one-sided case, the twosided case led to higher levels of generalization (Figure 3c).

**Fig. 4.**
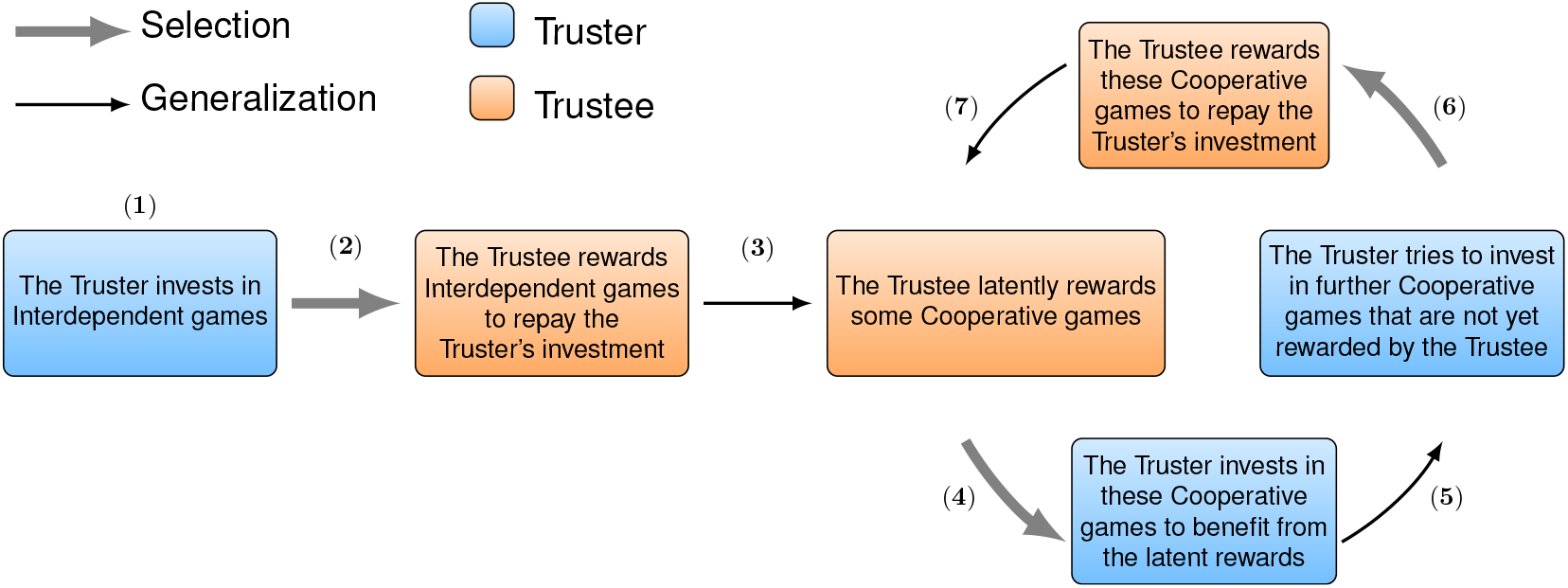
Evolutionary steps leading to the emergence of cooperation. (1) The truster is under selection to invest in Interdependent games. (2) As a result, the trustee is under selection to reward the truster in Interdependent games, when the automatic reward happens to be insufficient on its own. (3) As a byproduct, the trustee has a latent ability to reward some Cooperative games as well. (4) The truster is under selection to take advantage of latent rewards by investing in these Cooperative games. (5) Although the truster is under selection to avoid investing in other Cooperative games, in practice it is difficult to avoid every false positive. Hence the truster invests in some Cooperative games that the trustee does not yet reward. These false positive play the same role as latent rewards, only the other way around. Their existence generates a selection pressure on the trustee to reward the games where the truster happens to invest. (6) As a result, the set of games for which rewarding is under selection increases and generalization becomes stronger on the trustee side. (7) Since there are more and more Cooperative games to invest in, the truster makes further false positives.

**Fig. 5.**
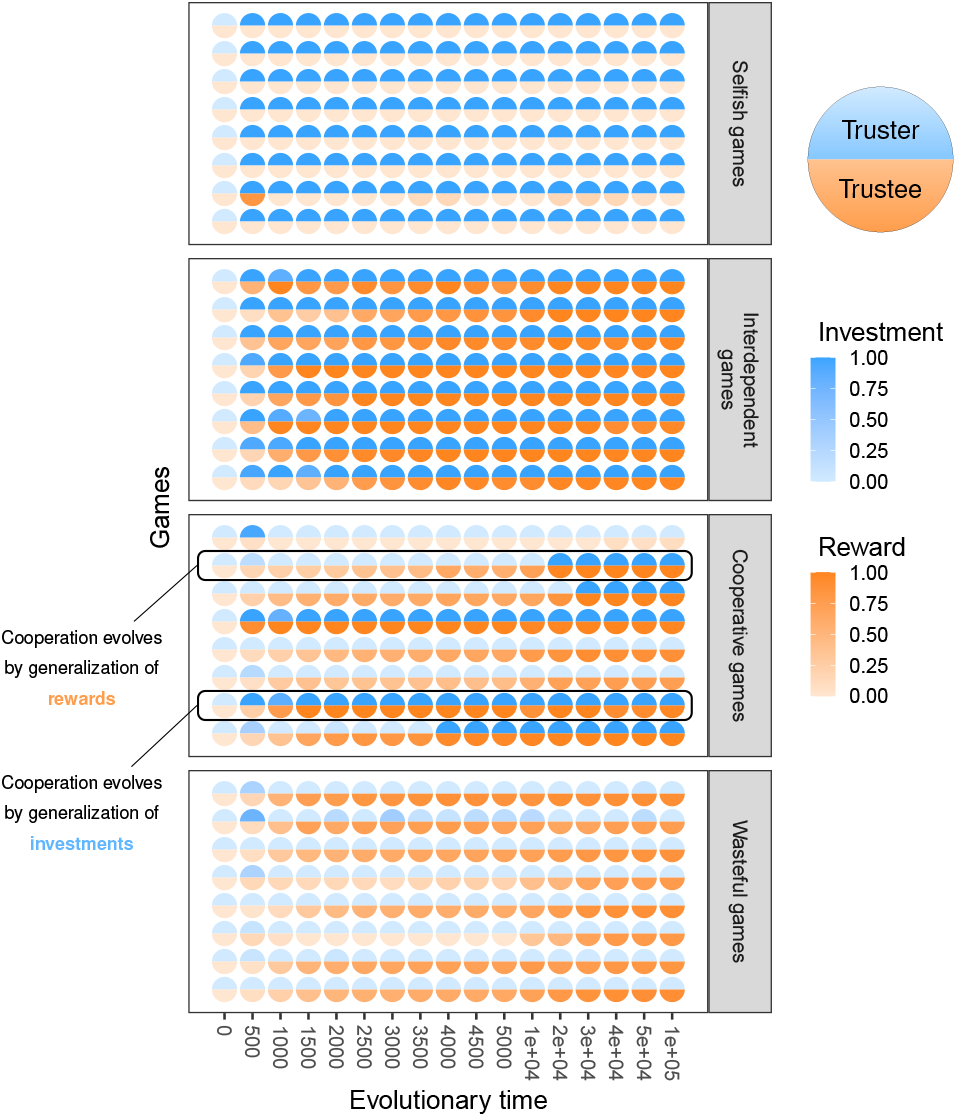
The joint evolution of cooperation and rewards outlined in one simulation example. We show 8 games of each type, chosen at random. For interdependent games, we only show the active reward, *r*, that the trustees offer when the automatic reward turns out to be absent (i.e. when *r_a_* = 0). Note that, for each game, we show a rescaled value *r/b* for the reward so that this value is 1 when the trustees provide a sufficient reward. See *Methods* for more details and for parameters.

In addition to the number of interdependent games and the cognitive complexity of the task, the evolution of cooperation also depended on the relative importance of Interdependent and Cooperative games in individuals’ ecology. The greater the role that cooperation plays overall in the ecology of individuals, the easier is evolutionary generalization (Fig. 6).

**Fig. 6.**
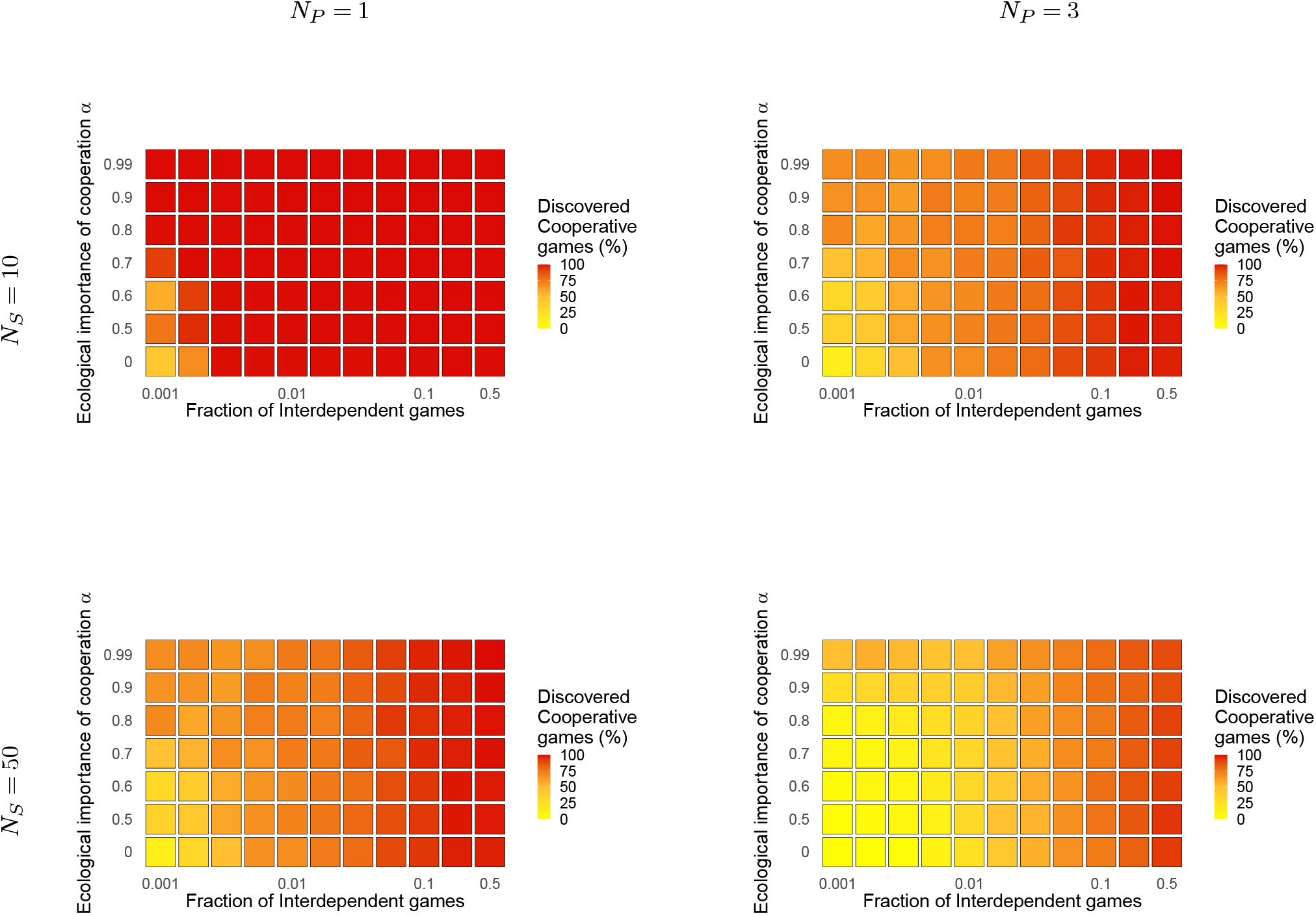
The ecological importance of cooperation, and evolutionary generalization. We show the equilibrium percentage of Cooperative games which are both played by the trusters and rewarded bythe trustees in function of (i) the fraction of Interdependent games and (ii) the relative ecological importance of cooperation *α* (relative importance of Interdependent and Cooperative games on fitness compared to Selfish and Wasteful games, see *Methods*). The case *α* = 0 corresponds to the results in Figure 3c and 3d. Results are shown fortwo values of the number *N_S_* of spurious features and for two values of the number *N_P_* of inputs per payoff parameter. Each point is the average value over 30 simulations.

## Discussion

Evolution by natural selection requires that it is always the same individual who expresses a trait and benefits from it. Each time this constraint has been partially lifted in the course of evolution –through genetic kinship or through the evolution of co-replication– a major transition took place, as this broadened the range of adaptive traits that could evolve (55, 56). In principle, reciprocity should be the most powerful means of lifting this constraint, as it can bring about the evolution of any trait by which an individual provides a benefit to any other, on the sole condition that the latter is able to offer some benefit back.

What makes this mechanism so powerful also has an evolutionary downside, however. Reciprocal cooperation faces a chicken and egg problem. The ability to reward others’ cooperation is an adaptation to the possibility of others cooperating, but this possibility is itself an adaptation to the fact that cooperation is rewarded. This probably explains why, despite its potency, reciprocal cooperation plays a much more limited role in the major transitions, than other mechanisms of cooperation (57).

In this article, we have sought to understand what evolutionary mechanisms can account for the emergence of reciprocal cooperation in spite of this problem. We have shown that this emergence is possible under the combined effect of two mechanisms: (i) a selective pressure resulting from specific ecological situations that play the role of triggers and (ii) an evolutionary mechanism analogue to generalisation.

The triggering ecological situations are social interactions wherein individuals have a *partner-dependent* degree of common interest with others. Typical instances of these are collective actions where both partners have a stake. The benefit that each party derives from the collective action depends on the quality of the other. Given that several partners or, more generally, several different opportunities to participate in a collective action may be put in competition with each other, it is natural that some are profitable and some are not (58). This selects for an ability to cooperate conditionally on partner’s quality. In turn, this ability creates a selection pressure favouring individuals of high quality, that is individuals who give others a reward large enough to compensate for the cost of their cooperation. In evolutionary equilibrium, therefore, individuals cooperate reciprocally, without any chicken and egg problem.

On the other hand, these reciprocal mechanisms have no reason to operate outside the specific interactions in which they have evolved in the first place. This explains why the overwhelming majority, and perhaps arguably all instances of reciprocal cooperation documented outside humans actually correspond to situations where individuals have some degree of shared interest in cooperating (46, 59). Reciprocal cooperation leads to a quantitative escalation of forms of cooperation that were already advantageous at the outset, in the absence of reciprocation. It does not allow the emergence of *novel* forms of cooperation that obligatorily need to be based on reciprocity to be adaptive. Put differently, as several scholars have noted, cooperation in nature is more frequently found in snowdrift games – where individuals have partially overlapping interests – than in prisoners’ dilemmas –where individuals have no shared interest (59–61).

Hence, reciprocal cooperation does not truly deliver on its promise to yield cooperation in all situations where individuals can generate benefits for each other. Its scope is considerably more limited in practice. Should the individuals of a species be able to overcome this constraint, they would dramatically broaden the range of benefits they are able to collect.

In this paper, we have shown that this is possible if individuals’ ecology encompasses not just a few triggering situations but a large number of them in a wide variety of areas of their lives. In this case, selection favors reciprocal mechanisms that can generalize beyond the situations for which they were originally selected. Individuals are not only good hunting partners or good nectar producers to attract pollinators, rather they are able *in general* to assess the cost paid by others in cooperating, and ensure that they are repaid by cooperating the right amount in return. Conversely, individuals have no reason to cooperate only when they have an immediate interest in doing so. Rather, they seek to cooperate whenever they are in a position to give a benefit to someone else, provided she has the opportunity to give a benefit in return.

To our current knowledge, the human species is the only one in which general mechanisms of this kind have evolved. Our fairness-based cooperation consists in calculating the costs paid by others in cooperating, whatever they may be, and reimbursing them by cooperating back in any way we can (62–64). This allows us to generate positive effects on each other in a extraordinarily wide range of situations, compared to other species in which reciprocal cooperation remains limited to the few situations where, for contingent reasons, the bootstrapping problem could be overcome.

## Methods

### Initialization

We run simulations coded into Python. In each simulation, a set of games is generated which contains 500 Selfish games, 250 Cooperative games, 250 Wasteful games, and a number of Interdependent games which varies between 1 and 2^10^ depending on the simulation. For each game, we generate random payoff parameters (*c* ∈ [−1,1], *b* ∈ [0,1] and *r_a_* ∈ [0,1] for Interdependent games) under the conditions provided in section *Games of life*.

In real life, individuals do not have a straightforward knowledge of the costs and benefits of the interaction in which they are involved. Rather, they have to make decisions based on various sensory inputs from the environment. Some inputs carry partial information about the payoff parameters, while others are just spurious features which happen to be correlated with the current game. In our simulations, for each payoff parameter, players do not receive its actual value *y* ∈ {*b,c,r_a_*}, but rather a number *N_P_* of inputs 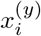 from which they can infer the true payoff parameter value *y*. At the beginning of each simulation, we generate three functions (for the three payoff parameter *b*, *c* and *r_a_*) that the players have to “learn” over evolutionary time if they want to base their decisions on the payoff parameters values:

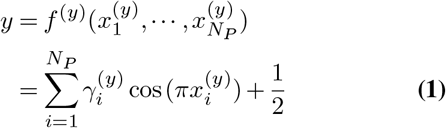

Parameters 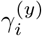 are randomly chosen, the only restriction being that there always exist a set 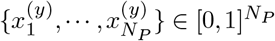 which satisfies (1) for all the possible values of the payoff parameter *y*. The number *N_P_* of inputs per payoff parameter determines the complexity of the task, which consists of inferring the “true” payoff parameter value from the sensory inputs. Additionally, for each game, we randomly choose *N_S_* spurious inputs on which the players can also base their decisions. In total, the truster receives 2*N_P_* + *N_S_* inputs and the trustee receives 3*N_P_* + *N_S_* inputs, since the truster does not have access to the automatic reward *r_a_* when he decides to “pay to see” the possible reward (see Figure 1).

Two Artificial Neural Networks (ANN), the truster and the trustee, are generated with a single latent layer of 10 neurons and with randomly chosen synaptic weights. For each vector of inputs (payoff inputs and spurious features of a single game), the truster’s output is a probability *p* ∈ [0,1] to “pay to see” the reward for this game. Note that we only explicitely model the first decision of the truster, which is to decide whether or not to “pay to see” the trustee’s reward (Figure 1). We assume that the second decision, which is to invest or to decline after monitoring the reward, is optimized by natural selection and that the truster invests iff *r_a_* + *r* − *c*. The trustee’s output is the reward value *r* for this game.

### Pre-selection phase

It can happen that, by chance, randomly generated ANNs monitor rewards or provide a large enough reward to the truster in some Cooperative games (*r > c*). This would be cases of fortuitous evolution of cooperation not driven by generalization, which is precisely what we are interested in. To avoid these situations, we first train the ANNs to never monitor (*p* = 0) and to provide a null reward in every games (*r* = 0).

### Selection phase

We then train both ANNs to provide respectively the adaptive probability of “pay to see” and the adaptive reward in each game of the training set as explained in section *Latent rewards solve the bootstrapping of reciprocal cooperation*. Note that, the trustee’s ANN is not trained in some games: the games in which the truster never invests (*p* = 0).

### Gradient descent

Previous works have already studied the evolution of social behaviours with ANNs (Arak and Enquist 1993; Enquist and Arak 1993; Johnstone 1993; André and Nolfi 2016; Debove et al. 2017). These works use genetic algorithms inspired from evolution by natural selection. However, there is a formal equivalence between natural selection and simple forms of learning (Watson and Szathmáry 2016, Zador 2019; Hasson et al. 2020). For instance, many machine learning methods are based on gradient descent. This principle is also used in a class of models which have been extensively used to described the evolutionary process: adaptive dynamics (Hofbauer and Sigmund 1990; Geritz et al. 1998). In the case of natural selection, the organisms’ traits gradually evolve towards maximisation of fitness. In the case of gradient descent, the ANN’s synaptic weights are gradually modified according to the gradient of the error function which quantifies how accurate the ANN is. Both mechanisms lead to decision machines which provide the accurate output for every set of inputs that they have encountered.

In our simulations, we use a simple back-propagation algorithm, i.e. gradient descent applied to ANNs (Goodfellow et al. 2016). More precisely, we use the mini-batch gradient descent with a batch size of 50 and a learning rate of 0.2. At each iteration, we compute the outputs of the truster (*p*) and the trustee (*r*) in 50 randomly drawn games. From these, we infer their adaptive values *p\** and *r\** in each games (see section *The truster’s evolutionarily stable strategy*) and we perform a single step of back-propagation of both ANNs to update the synaptic weights in the direction that maximizes the fitness, i.e. that minimizes the difference between the actual output and its adaptive value. We use the following error function for the truster

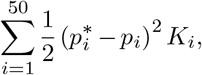

and for the trustee

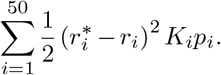

Note that the difference is multiplied by a factor to take into account the fact that selection is stronger in some games than in others. This method is known as cost-sensitive learning (Kukar & Kononenko 1998). During the updating step of back-propagation, each game contributes more or less in the error function depending on their type. Namely, the relative ecological importance of cooperation *α* is taken into account by setting *K_i_* = 1 for Interdependent and Cooperative games and *K_i_* = 1 − *α* for Selfish and Wasteful games. If *α* = 0, the strength of selection is the same in all games. On the opposite, if *α* = 1, only Interdependent and Cooperative games have an effect on the fitness. We always assume that *α* = 0, except for results shown in Figure 6. In addition, we assume that selection on the trustee to provide a reward in a game *i* linearly increases with the probability *p_i_* that the truster “pays to see” the reward in this game.

For each simulation, we perform 10^3^ minibatch steps for the pre-selection phase and 10^6^ for the selection. The equilibrium values shown in Figures 3 and 6 are average values over the last 5 × 10^4^ steps where we make sure that the algorithm has converged.

### The truster’s evolutionarily stable strategy

The truster must decide, for every game, whether he wants to decline, or play conditionally on the trustee’s quality. We have modelled this decision with a probability *p* ∈ [0,1] to “pay to see”.

Monitoring the trustee in a given game can only be worth it if the sign of the truster’s net payoff in this game is uncertain (that is, if *r_a_* + *r* − *c* is sometimes positive and sometimes negative in the same game, depending on the trustee’s quality). Otherwise, it is always better for the truster to spare the monitoring cost, and play or decline the game unconditionally. If the sign of *r_a_* + *r* − *c* is uncertain, on the other hand, the truster’s optimal strategy is less straightforward. In the sake of simplicity, throughout the paper, we assume that monitoring is always worth it in this case. That is, we assume that interactions are long (or more generally important in terms of payoff), such that the monitoring cost *m* is always lower than both the expected benefit of beneficial interactions and the expected cost of costly interactions.

In consequence, the evolutionarily stable strategy (ESS) of the truster is the following:

1. In all games where *r* + *r_a_* − *c* is always negative, the truster must decline (*p\** = 0).
2. In all games where the sign of *r* + *r_a_* − *c* is uncertain, the truster must pay *m* to monitor the trustee’s quality (*p*\* = 1) and then play the game iff *r* + *r_a_* − *c*> 0

### The trustee’s evolutionarily stable strategy

The trustee should actively produce a reward only when it is worth it, that is only when the truster takes her reward into account to make a decision. Accordingly, the evolutionarily stable strategy of the trustee is the following:

1. In all games where the truster will play conditionally on the sign of *r* + *r_a_ − c*, the trustee should offer a reward *r\** = *c* − *r_a_* + *ϵ*, where *ϵ*> 0 is just sufficient to incentivize the truster to play the game (see 65).
2. In all games where the truster will, anyway, decline unconditionally, the trustee’s decision is neutral because it never actually occurs. It constitutes a “latent” facet of the trustee’s strategy.

## Notes

### Competing Interest Statement

The authors have declared no competing interest.

## References

1. George Christopher Williams. Adaptation and Natural Selection: A Critique of Some Current Evolutionary Thought. Princeton Science Library, 1966.

2. W D Hamilton. The genetical evolution of social behaviour, I & II. J Theor Biol, 7:1–52, 1964.

3. S A West, A S Griffin, and A Gardner. Social semantics: Altruism, cooperation, mutualism, strong reciprocity and group selection. Journal of Evolutionary Biology, 20(2):415–432, 2007. ISSN 1010061X. doi: 10.1111/j.1420-9101.2006.01258.x.

4. R L Trivers. The evolution of reciprocal altruism. Quarterly Review of Biology, 46:35–57, 1971.

5. Robert Axelrod and William D Hamilton. The evolution of cooperation. Science, 211(4489):1390–1396, 1981. ISSN 08893047. doi: 10.1007/BF01539342.

6. D Fudenberg and E Maskin. The Folk Theorem in Repeated Games with Discounting or with Incomplete Information. Econometrica, 54(3):533–554, 1986.

7. K Binmore and L Samuelson. Evolutionary stability in repeated games played by finite automata. Journal of Economic Theory, 57:278–305, 1992.

8. MA Nowak and K Sigmund. Tit for tat in heterogeneous populations. Nature, 355:250–253, 1992.

9. Robert Boyd and Peter J. Richerson. Punishment allows the evolution of cooperation anything else) in sizable groups. Ethology and Sociobiology, 13(3):171–195, 1992. ISSN 01623095. doi: 10.1016/0162-3095(92)90032-Y.

10. M Nowak and K Sigmund. A Strategy of Win Stay, Lose Shift That Outperforms Tit-for-Tat in the Prisoners-Dilemma Game. Nature, 364(6432):56–58,1993.

11. MA Nowak and K Sigmund. Evolution of indirect reciprocity by image scoring. Nature, 393 (6685):573–577, 1998.

12. G Roberts and T N Sherratt. Development of cooperative relationships through increasing investment. Nature, 394(6689):175–179, 1998.

13. Olof Leimar and Peter Hammerstein. Evolution of cooperation through indirect reciprocity, 2001. ISSN 14712970.

14. C Aktipis. Know when to walk away: contingent movement and the evolution of cooperation. Journal of Theoretical Biology, 231(2):249–260, 2004.

15. John M Mcnamara, Zoltan Barta, Lutz Fromhage, and Alasdair I Houston. The coevolution of choosiness and cooperation. 451(January):189–192, 2008. doi: 10.1038/nature06455.

16. Pat Barclay. Competitive helping increases with the size of biological markets and invades defection. Journal of Theoretical Biology, 281(1):47–55, jul 2011. ISSN 00225193. doi: 10.1016/j.jtbi.2011.04.023.

17. J. Arvid Ågren, Nicholas G. Davies, and Kevin R. Foster. Enforcement is central to the evolution of cooperation, 2019. ISSN 2397334X.

18. Jean-Baptiste André. Mechanistic constraints and the unlikely evolution of reciprocal cooperation. Journal of Evolutionary Biology, 27(4):784–795, mar 2014. ISSN 14209101. doi: 10.1111/jeb.12351.

19. Jean-Baptiste André and Stefano Nolfi. Evolutionary robotics simulations help explain why reciprocity is rare in nature. Scientific Reports, in press, 2016.

20. R Dawkins and J R Krebs. Animal signals: information or manipulation. Behavioural ecology: An evolutionary approach, pages 282–309, 1978.

21. J R Krebs, R Dawkins, and Others. Animal signals: mind-reading and manipulation. Behavioural Ecology: an evolutionary approach, 2:380–402, 1984.

22. Thomas C. Scott-Phillips, Richard A. Blythe, Andy Gardner, and Stuart A. West. How do communication systems emerge? Proceedings of the Royal Society B: Biological Sciences, 279(1735):1943–1949, 2012. ISSN 14712970. doi: 10.1098/rspb.2011.2181.

23. Michael Tomasello, Alicia P. Melis, Claudio Tennie, Emily Wyman, and Esther Herrmann. Two key steps in the evolution of human cooperation: The interdependence Hypothesis. Current Anthropology, 53(6):673–692, dec 2012. ISSN 00113204. doi: 10.1086/668207.

24. Dan Sperber and Nicolas Baumard. Moral Reputation: An Evolutionary and Cognitive Perspective. Mind and Language, 27(5):495–518, 2012. ISSN 02681064. doi: 10.1111/mila.12000.

25. Yaser S. Abu-Mostafa, Malik. Magdon-Ismail, and H T Lin. Learning from Data: A Short Course. 2012. ISBN 1600490069.

26. A. Arak and M. Enquist. Hidden preferences and the evolution of signals. Philosophical Transactions of the Royal Society of London. Series B: Biological Sciences, 340(1292):207–213, may 1993. ISSN 0962-8436. doi: 10.1098/rstb.1993.0059.

27. Magnus Enquist and Anthony Arak. Selection of exaggerated male traits by female aesthetic senses. Nature, 361(6411):446–448, 1993. ISSN 0028-0836. doi: 10.1038/361446a0.

28. Rufus A. Johnstone. Female preference for symmetrical males as a by-product of selection for mate recognition. Nature, 372(6502):172–175, 1994. ISSN 00280836. doi: 10.1038/372172a0.

29. M. Enquist and R. A. Johnstone. Generalization and the evolution of symmetry preferences. Proceedings of the Royal Society B: Biological Sciences, 1997. ISSN 14712970. doi: 10.1098/rspb.1997.0186.

30. Magnus Enquist, Anthony Arak, Stefano Ghirlanda, and Carl-Adam Wachtmeister. Spectacular phenomena and limits to rationality in genetic and cultural evolution. Philosophical transactions of the Royal Society of London. Series B, Biological sciences, 357(1427):1585–94, nov 2002. ISSN 0962-8436. doi: 10.1098/rstb.2002.1067.

31. Stefano Ghirlanda and Magnus Enquist. A century of generalization, 2003. ISSN 00033472.

32. Richard A Watson and Eörs Szathmáry. How Can Evolution Learn? Trends in Ecology & Evolution, 31(2):1–11, 2015. ISSN 01695347. doi: 10.1016/j.tree.2015.11.009.

33. Kostas Kouvaris, Jeff Clune, Loizos Kounios, Markus Brede, and Richard A. Watson. How evolution learns to generalise: Using the principles of learning theory to understand the evolution of developmental organisation. PLoS Computational Biology, 2017. ISSN 15537358. doi: 10.1371/journal.pcbi.1005358.

34. Uri Hasson, Samuel A. Nastase, and Ariel Goldstein. Direct Fit to Nature: An Evolutionary Perspective on Biological and Artificial Neural Networks, 2020. ISSN 10974199.

35. Merav Parter, Nadav Kashtan, and Uri Alon. Facilitated variation: how evolution learns from past environments to generalize to new environments. PLoS computational biology, 4(11):e1000206, nov 2008. ISSN 1553-7358. doi: 10.1371/journal.pcbi.1000206.

36. Larry Samuelson. Analogies, adaptation, and anomalies. Journal of Economic Theory, 97(2):320–366, 2001.

37. Jenna Bednar and Scott Page. Can game (s) theory explain culture? {The} emergence of cultural behavior within multiple games. Rationality and Society, 19(1):65–97, 2007.

38. Jenna Bednar, Yan Chen, Tracy Xiao Liu, and Scott Page. Behavioral spillovers and cognitive load in multiple games: {An} experimental study. Games and Economic Behavior, 74(1):12–31, 2012.

39. Friederike Mengel. Learning across games. Games and Economic Behavior, 74(2):601–619, 2012.

40. Olof Leimar and Richard C RC Connor. By-product benefits, reciprocity, and pseudoreciprocity in mutualism. In Peter Hammerstein, editor, Genetic and Cultural Evolution of Cooperation, pages 203–222. The MIT Press, Cambridge MA, 2003. ISBN 0262083264.

41. R C Connor. Pseudo-reciprocity: investing in mutualism. Animal Behaviour, 34(5):1562–1566, 1986.

42. I Eshel and A Shaked. Partnership. Journal of theoretical biology, 208(4):457–74, feb 2001. ISSN 0022-5193. doi: 10.1006/jtbi.2000.2232.

43. Hanna Kokko and Rufus A Johnstone. The evolution of cooperative breeding through group augmentation. Proceedings of the Royal Society of London. Series B: Biological Sciences, 268(1463):187–196, 2001. doi: 10.1098/rspb.2000.1349.

44. G Roberts. Cooperation through interdependence. Animal Behaviour, 70(4):901–908,2005.

45. Tim Clutton-Brock. Cooperation between non-kin in animal societies. Nature, 462(7269):51–7, nov 2009. ISSN 1476-4687. doi: 10.1038/nature08366.

46. Olof Leimar and Peter Hammerstein. Cooperation for direct fitness benefits. Philosophical Transactions of the Royal Society B-Biological Sciences, 365(1553):2619–2626, 2010. doi: DOI10.1098/rstb.2010.0116.

47. Martin Heil, Marcia González-Teuber, Lars W Clement, Stefanie Kautz, Manfred Verhaagh, and Juan Carlos Silva Bueno. Divergent investment strategies of Acacia myrmecophytes and the coexistence of mutualists and exploiters. Proceedings of the National Academy of Sciences of the United States of America, 106(43):18091–18096, 2009. ISSN 00278424. doi: 10.1073/pnas.0904304106.

48. Graham H. Pyke. Plant-pollinator co-evolution: It’s time to reconnect with Optimal Foraging Theory and Evolutionarily Stable Strategies, 2016. ISSN 16180437.

49. D. P. Croft, R. James, P. O. R. Thomas, C. Hathaway, D. Mawdsley, K. N. Laland, and J. Krause. Social structure and co-operative interactions in a wild population of guppies (Poecilia reticulata). Behavioral Ecology and Sociobiology, 59(5):644–650, nov 2005. ISSN 0340-5443. doi: 10.1007/s00265-005-0091-y.

50. Foster Provost. Machine learning from imbalanced data sets 101. In Proceedings of the {AAAI}’2000 workshop on imbalanced data sets, pages 1–3, 2000.

51. Gary M Weiss and Foster Provost. The effect of class distribution on classifier learning: an empirical study. Rutgers Univ, 2001.

52. Nathalie Japkowicz and Shaju Stephen. The class imbalance problem: {A} systematic study. Intelligent data analysis, 6(5):429–449, 2002.

53. Maciej A Mazurowski, Piotr A Habas, Jacek M Zurada, Joseph Y Lo, Jay A Baker, and Georgia D Tourassi. Training neural network classifiers for medical decision making: {The} effects of imbalanced datasets on classification performance. Neural networks, 21(2-3):427–436, 2008.

54. Haibo He and Edwardo A Garcia. Learning from imbalanced data. IEEE Transactions on Knowledge & Data Engineering, (9):1263–1284, 2009.

55. J Maynard Smith and E Szathmary. The major transitions in evolution. Oxford University Press, Oxford, 1995.

56. Richard E. Michod. Darwinian Dynamics: Evolutionary Transitions in Fitness and Individuality, 2000. ISSN 00063568.

57. P Hammerstein. Why is reciprocity so rare in social animals? A protestant appeal. In P Hammerstein, editor, Genetic and cultural evolution of cooperation, pages 83–94. MIT Press, Cambridge, MA, 2003.

58. Ronald Noë and Peter Hammerstein. Biological markets: supply and demand determine the effect of partner choice in cooperation, mutualism and mating. Behavioral Ecology and Sociobiology, 35(1):1–11, 1994. ISSN 03405443. doi: 10.1007/BF00167053.

59. Redouan Bshary, Klaus Zuberbühler, and Carel P. Van Schaik. Why mutual helping in most natural systems is neither conflict-free nor based on maximal conflict. Philosophical Transactions of the Royal Society B: Biological Sciences, 371(1687), 2016. ISSN 14712970. doi: 10.1098/rstb.2015.0091.

60. N J Raihani and R Bshary. Resolving the iterated prisoner’s dilemma: theory and reality. Journal of Evolutionary Biology, 24(8):1628–39, aug 2011. ISSN 1420-9101. doi: 10.1111/j.1420-9101.2011.02307.x.

61. Redouan Bshary and Judith L. Bronstein. A General Scheme to Predict Partner Control Mechanisms in Pairwise Cooperative Interactions Between Unrelated Individuals. Ethology, 117(4):271–283, 2011. ISSN 01791613. doi: 10.1111/j.1439-0310.2011.01882.x.

62. Stéphane Debove, Nicolas Baumard, and Jean Baptiste André. On the evolutionary origins of equity. PLoS ONE, 12(3):5–7, 2017. ISSN 19326203. doi: 10.1371/journal.pone.0173636.

63. Jean-Baptiste André and Nicolas Baumard. Social opportunities and the evolution of fairness. Journal of Theoretical Biology, 289(1):128–135, nov 2011. ISSN 00225193. doi: 10.1016/j.jtbi.2011.07.031.

64. Nicolas Baumard, J.-B. Jean-Baptiste André, Dan Sperber, and Others. A mutualistic approach to morality: The evolution of fairness by partner choice. Behavioral and Brain Sciences, 36(1):59–122, 2013. ISSN 14691825. doi: 10.1017/S0140525X11002202.

65. Ken Binmore, Avner Shaked, and John Sutton. An outside option experiment. Quarterly Journal of Economics, 104(4):753–770, 1989. ISSN 15314650. doi: 10.2307/2937866.

